# Embryonic lympho-epithelial cell interactions play an essential role in the establishment of adult T cell tolerance

**DOI:** 10.64898/2026.01.19.700349

**Authors:** G. Nogueira, A. Chervova, F. Soares-da-Silva, A. Sergé, M. Irla, Cabe C. Moraes, A. Gaudin, P. Ferreirinha, A. Bandeira, NL. Alves, P. Pereira, A. Cumano

**Author notes:** Corresponding author: Ana Cumano Institut Pasteur, 25 Rue du Dr. Roux 75724 Paris, CEDEX15, France.

## Abstract

Thymic seeding progenitors (TSP) colonize the embryonic thymus in two successive waves, with the first wave uniquely contributing to lymphoid tissue inducer and invariant γδ T cells that drive primordial medullary thymic epithelial cell (mTEC) maturation. Although recognized during thymic organogenesis, their long-term impact on T cell selection and immune tolerance remains unclear. We show that selective depletion of first-wave TSP alters thymic architecture and TEC composition. This results in reduced mTEC numbers, failure of medullary islets to expand and coalesce, altered dendritic cell composition, and delayed regulatory T cell development. Single-cell transcriptional analysis revealed a decrease in the most immature and expanding compartments and signs of accelerated maturation. It also revealed abnormal TEC differentiation trajectories, expansion of post-Aire and mimetic TEC subsets, and altered expression of tissue-restricted antigen-coding genes. Functionally, first-wave TSP depletion resulted in impaired thymic recovery after injury, and age-associated autoimmunity, with elevated anti-nuclear antibodies and lymphocytic infiltration in peripheral tissues. Thus, first-wave TSP imprint long-lasting control of thymic function and immune tolerance.

**One sentence summary:** Early embryonic thymic seeding progenitors program thymic epithelial maturation and regulatory T cell development, ensuring long-term immune tolerance and protection from age-associated autoimmunity.

## Introduction

Human aging and cytotoxic therapies are associated with declining thymic function, resulting in delayed T-cell reconstitution, increased susceptibility to infection, autoimmunity, and cancer (reviewed in^1–3^). Thymic epithelial cells (TEC) are central to T cell development and tolerance: cortical TEC (cTEC) mediate positive selection, whereas medullary TEC (mTEC), together with dendritic cells (DC), eliminate autoreactive clones and promote regulatory T (Treg) cell differentiation^4–6^. This process depends on the autoimmune regulator Aire, which drives promiscuous expression of peripheral tissue-restricted antigens (TRA) in a subset of mTEC^7^. Aire expression during the perinatal period is necessary and sufficient to establish long-term immune tolerance^8^. Recent studies have revealed mTEC heterogeneous “mimetic” subsets that adopt coordinated peripheral tissue–like transcriptional programs^9,10^.

TEC differentiation is initiated during embryogenesis and relies on reciprocal interactions with hematopoietic cells, including lymphoid tissue inducer (LTi), invariant γδ, and CD4⁺ T cells^11–13^. Two waves of thymus-seeding progenitors (TSP) colonize the embryonic thymus; the first wave, unlike subsequent waves, generates LTi and invariant γδ T cells^12,14^, mediating mTEC maturation and Aire induction^11^. Transient blockade of IL-7R signaling between embryonic days (E)10–14 selectively depletes first-wave TSP and reduces LTi cells, invariant γδ T cells, and mature mTEC at birth^12^.

Whether the first wave of TSP exerts a lasting, non-redundant role in shaping thymic architecture, differentiation and regenerative capacity, ensuring tolerance, remains unresolved. Here, we show that selective depletion of first-wave TSP leads to persistent defects in adult thymic epithelial organization and composition. Single-cell transcriptomics revealed altered TEC differentiation trajectories, with accumulation of post-Aire and mimetic mTEC subsets and dysregulated Aire-dependent and -independent TRA expression. These alterations impaired thymic regenerative capacity after injury and compromised central tolerance, resulting in elevated anti-nuclear antibodies (ANA) and tissue-specific immune infiltration in aged mice. Our findings establish first-wave TSP progeny as indispensable architects of thymic epithelial maturation and lifelong immune tolerance.

## Results

### Depletion of the first TSP wave affects TEC and Treg cell numbers, but not those of conventional T cells after the neonatal period

We analyzed thymic epithelial and hematopoietic compartments in offspring from pregnant females treated with anti-IL7Rα antibody between E10.5–E14.5 (anti-IL7Rα mice) at multiple postnatal time points (**Fig. 1A**; **Supplementary Fig. 1–3**). TEC were identified as CD45⁻EpCAM⁺ cells and subdivided into cTEC (Ly51⁺) and mTEC (UEA-1⁺), with mature mTEC defined as MHC-II^high^ CD80⁺ cells, including the Aire⁺ subset^15^ (**Supplementary Fig. 1A**).

**Figure 1.**
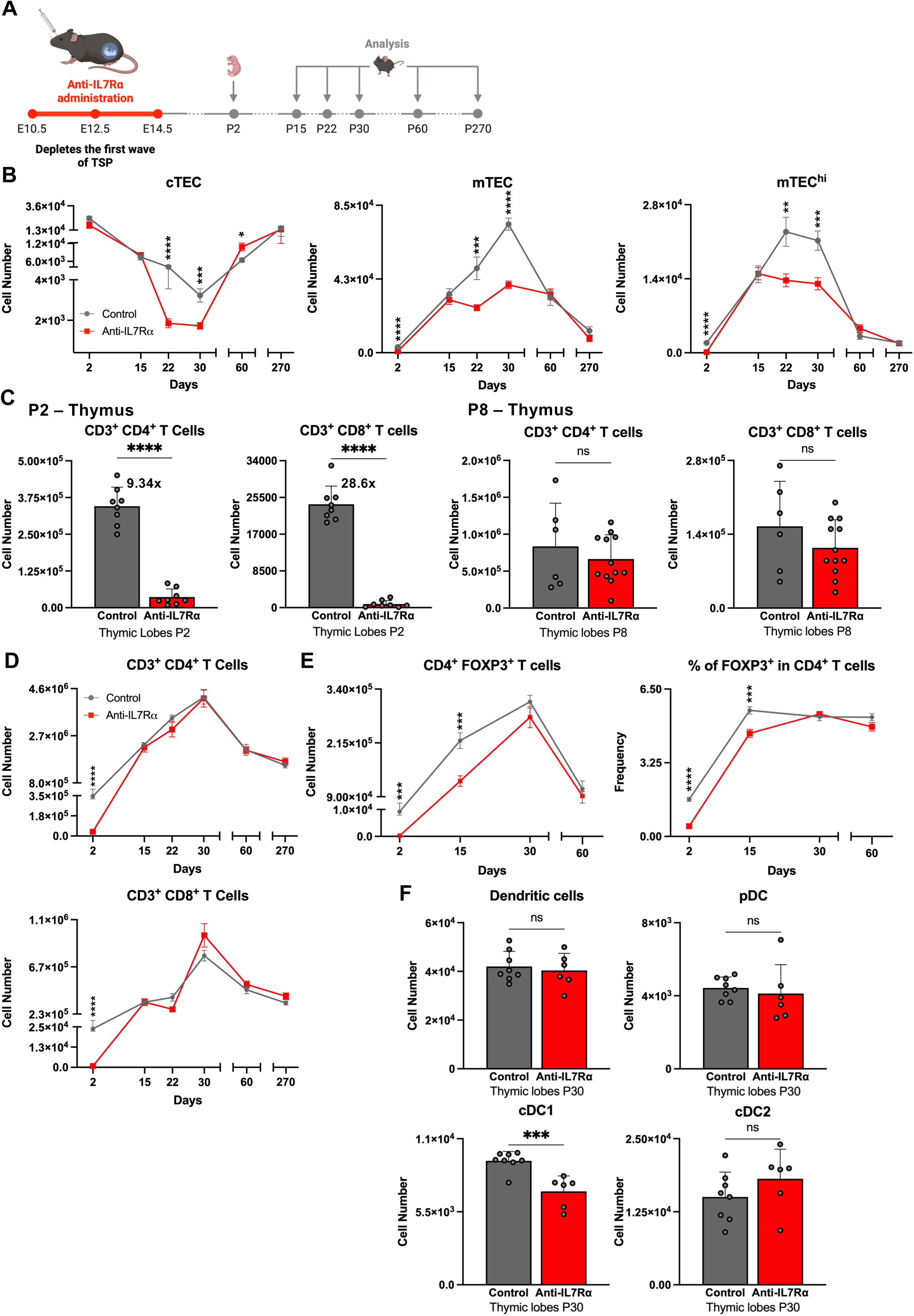
The first wave of TSP is indispensable for TEC maturation, Treg cell development, and DC balance in the thymus. **(A)** Experimental setup: pregnant C57BL/6J females were injected with anti-IL7Rα antibody at E10.5, E12.5, and E14.5; offspring were analyzed between postnatal day 2 (P2) and P270. **(B)** Absolute numbers of TEC subsets: cTEC (left), mTEC (middle) and mTEC^hi^ mTEC (right). **(C)** Total numbers of CD3^+^ CD4^+^ and CD8^+^ thymic T cells at P2 (left panels) and at P8 (right panels). The numbers above the bar graphs represent the fold change between the values of the two experimental groups. **(D)** Absolute numbers of CD3^+^ CD4^+^ (top) and CD8^+^ (bottom) T cells in the thymus between P2 and P270. (**E)** Absolute numbers (left) and frequency (right) of FOXP3⁺ cells within CD4⁺ T cells in the thymus between P2 and P60. **(F)** Absolute numbers of total DC, pDC, cDC1, and cDC2 in thymic lobes at P30. Each dot represents an individual mouse; control (grey) and anti-IL7Rα (red). Data are presented as the mean ± SEM (B, D, E) or SD (C and F) of ≥ 2 independent experiments. ns, not significant. *p < 0.05, **p < 0.01, ***p < 0.001, ****p < 0.0001 (Unpaired T-test).

In control mice, total TEC numbers increased after postnatal day (P)2 and reached a maximal number at P30, (**Fig. 1B**; **Supplementary Fig. 1B**), consistent with previous reports^16,17^. This expansion was driven primarily by mTEC, including mTEC^hi^, whereas the cTEC compartment – containing most immature TEC and putative bipotent progenitors^18,19^ – declined sharply over the same period (**Fig. 1B**; **Supplementary Fig. 1B**). In contrast, anti-IL7Rα mice exhibited significantly reduced numbers of total mTEC and mTEC^hi^ at P2 that further increased until P15 but failed to expand further thereafter (**Fig. 1B**; **Supplementary Fig. 1B**). The cTEC compartment in treated mice declined more rapidly and reached lower absolute numbers than in controls. After P30, mTEC numbers in anti-IL7Rα mice remained low and relatively stable from P15 to P60 declining further thereafter, whereas in control mice they declined after P30 as expected (**Fig. 1B**; **Supplementary Fig. 1B**). In both groups, mTEC loss was accompanied by a rebound in cTEC numbers, resulting in comparable mTEC/cTEC ratios throughout postnatal life (**Supplementary Fig. 1C**). These data indicate that mTEC in anti-IL7Rα mice display a restricted capacity for postnatal expansion.

We previously showed that anti-IL7Rα treatment impairs the first wave of TSP, resulting at P2 in reduced numbers of SP4 thymocytes, invariant Vγ5Vδ1 T cells, and LTi cells, but normal numbers of second-wave-derived DN thymocytes^12^. A refined analysis confirmed the reduction in mature SP4 (∼10-fold) with an even more pronounced reduction in SP8 (∼29-fold) thymocytes at P2 (**Fig. 1C–D**) and a FOXP3⁺ compartment selectively affected in both frequency (3.6-fold) and absolute number (33-fold) (**Fig. 1E**; **Supplementary Fig. 2A**).

To examine postnatal T cell differentiation dynamics, we analyzed Rag2-GFP reporter mice (**Fig. 1C**; **Supplementary Fig. 2B–D**). At P8, SP4 and SP8 thymocyte numbers in anti-IL7Rα mice had reached levels comparable to controls and displayed similar proportions of GFP⁺ cells, indicating normal ongoing thymopoiesis (**Fig. 1C**; **Supplementary Fig. 2B**). Consequently, the expansion of mature thymocytes between P2 and P8 was markedly accelerated in anti-IL7Rα mice (18.1-fold for SP4 and 136.7-fold for SP8), compared to controls (2.4-fold and 6.7-fold, respectively) (**Supplementary Fig. 2C**). Peripheral T cell seeding remained delayed at P8, with significantly fewer splenic CD4⁺ and CD8⁺ T cells in anti-IL7Rα mice (**Supplementary Fig. 2D**). However, GFP expression among peripheral CD4⁺ T cells was comparable between groups, with only a modest increase among CD8⁺ T cells in anti-IL7Rα mice (**Supplementary Fig. 2D**). After P8, thymic SP4 and SP8 cell numbers remained similar between groups (**Fig. 1D**). In contrast, Treg cell numbers and frequency among SP4 thymocytes remained reduced in anti-IL7Rα mice and only normalized by P30 (**Fig. 1E**), indicating impaired Treg differentiation.

At P30, plasmacytoid DC and cDC2 compartments were comparable between groups, whereas conventional dendritic cell type 1 (cDC1) numbers, implicated in negative selection and Treg induction^5^ were significantly reduced in anti-IL7Rα mice (**Fig. 1F**; **Supplementary Fig. 3**). As cDC1 are generated intrathymically^20^, this selective deficit suggests a non-redundant contribution of first-wave TSP to the development of this subset.

In summary, depletion of the first TSP wave leads to a exacerbated expansion of conventional SP4 and SP8 thymocytes during early postnatal life, while Treg cell development is selectively delayed. Normal thymocyte numbers after P8 contrast with the limited postnatal expansion of the TEC compartment between P15 and P30, resulting in an altered thymocyte-to-TEC balance and impaired Treg cell differentiation.

### Depletion of first-wave TSP disrupts thymic medullary organization

Initially arising as mTEC islets of clonal origin, these units progressively coalesce during thymic development to form a large and unified medulla in the adult thymus^21^. Given the critical role of primordial lymphoepithelial interactions in thymic medulla development^11^ and the low TEC-to-lymphocyte ratio in anti-IL7Rα mice we performed confocal microscopy of thymic sections and analyzed the spatial organization of the thymic medulla. We found a higher number of small medullary regions and a conspicuous absence of large, well-organized medullas as compared to control counterparts (**Fig. 2A–C**). These anatomical alterations persisted at P60, although both groups had similar numbers of mTEC (**Fig. 2D**; **Supplementary Fig. 4**). Three-dimensional reconstruction of the thymus architecture confirmed that anti-IL7Rα thymi harbored a greater number of peripheral medullary islets (111 vs. 64 per lobe in controls) (**Fig. 2E–F**; **Supplementary Video 1-2**). Together, these data indicate that thymocytes generated by the first wave are essential to induce a normal medullary organization, highlighting the non-redundant contribution of this TSP wave to thymic development and function.

**Figure 2.**
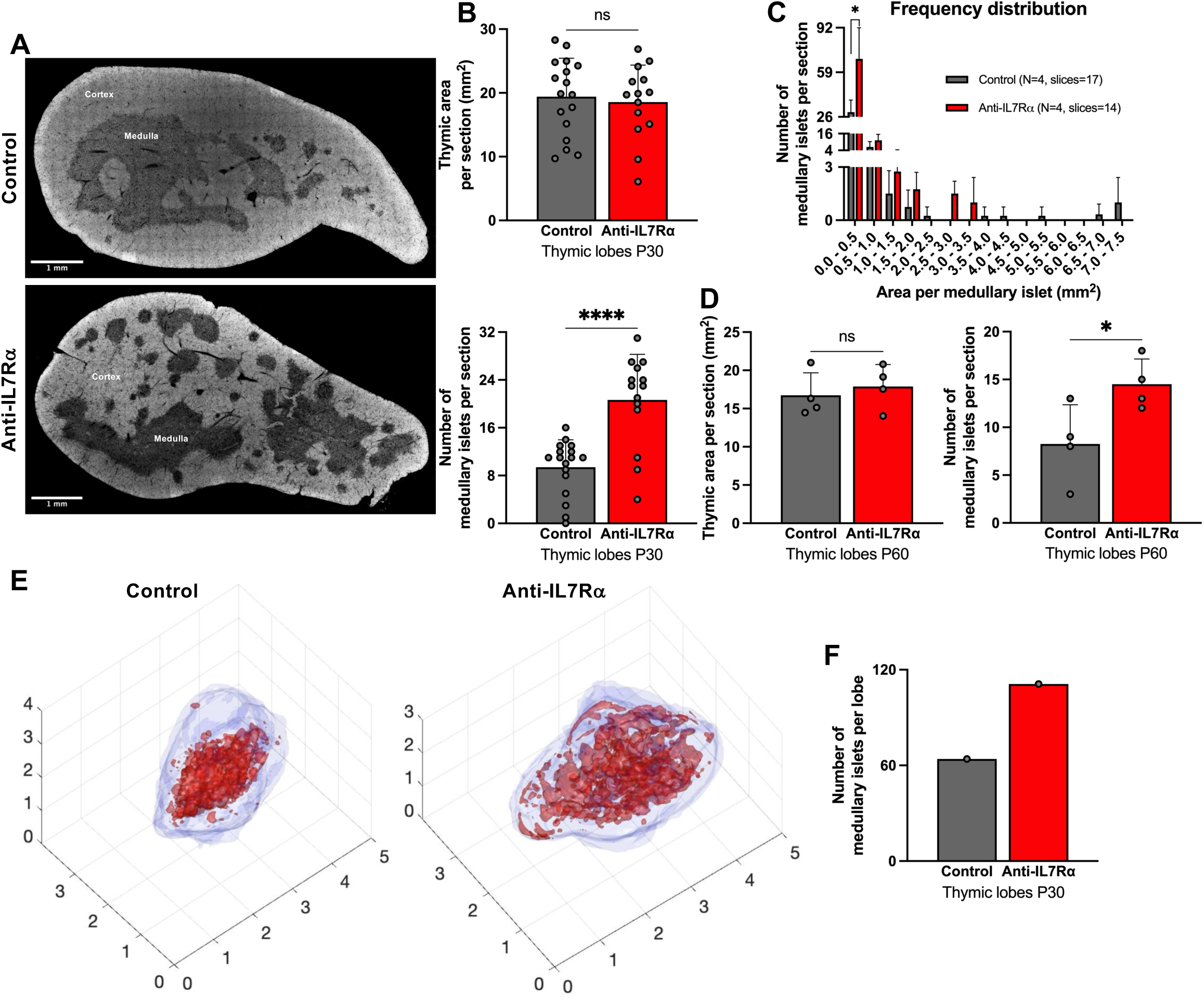
Long-term disruption of thymic medullary organization following depletion of first-wave TSP. **(A)** Representative DAPI-stained thymic sections at P30. **(B)** Quantification of thymic areas (top) and number of medullary islets per section (bottom) at P30. **(C)** Frequency distribution of medullary islet size in control and anti-IL7Rα mice. **(D)** Thymic area (left) and number of medullary islets (right) per section at P60. **(E)** 3D reconstructions of thymic lobes from control (left) and anti-IL7Rα (right) P30 mice. **(F)** Quantification of medullary islet number per thymic lobe. Control (grey) and anti-IL7Rα (red). Data are presented as the mean ± SD. ns, not significant. *p < 0.05, **p < 0.01, ***p < 0.001, ****p < 0.0001 (Unpaired T-test).

### Depletion of the first-wave TSP alters TEC diversity and transcriptional programs

To determine whether depletion of first-wave TSP impacts TEC differentiation, we performed single-cell RNA sequencing (10x Genomics) on sorted EPCAM⁺CD45⁻ TEC from control and anti-IL7Rα mice at P30, when differences in TEC cellularity are maximal (**Supplementary Fig. 1A–B**). Unsupervised clustering of 19,446 control and 15,857 anti-IL7Rα TEC identified 14 transcriptionally distinct subsets in both groups (**Fig. 3A**), corresponding to previously described TEC populations^9,22,23,24^ and annotated using the combined classification of Baran-Gale et al. ^24^ and Michelson et al.^9^ (**Supplementary Table 2**).

**Figure 3.**
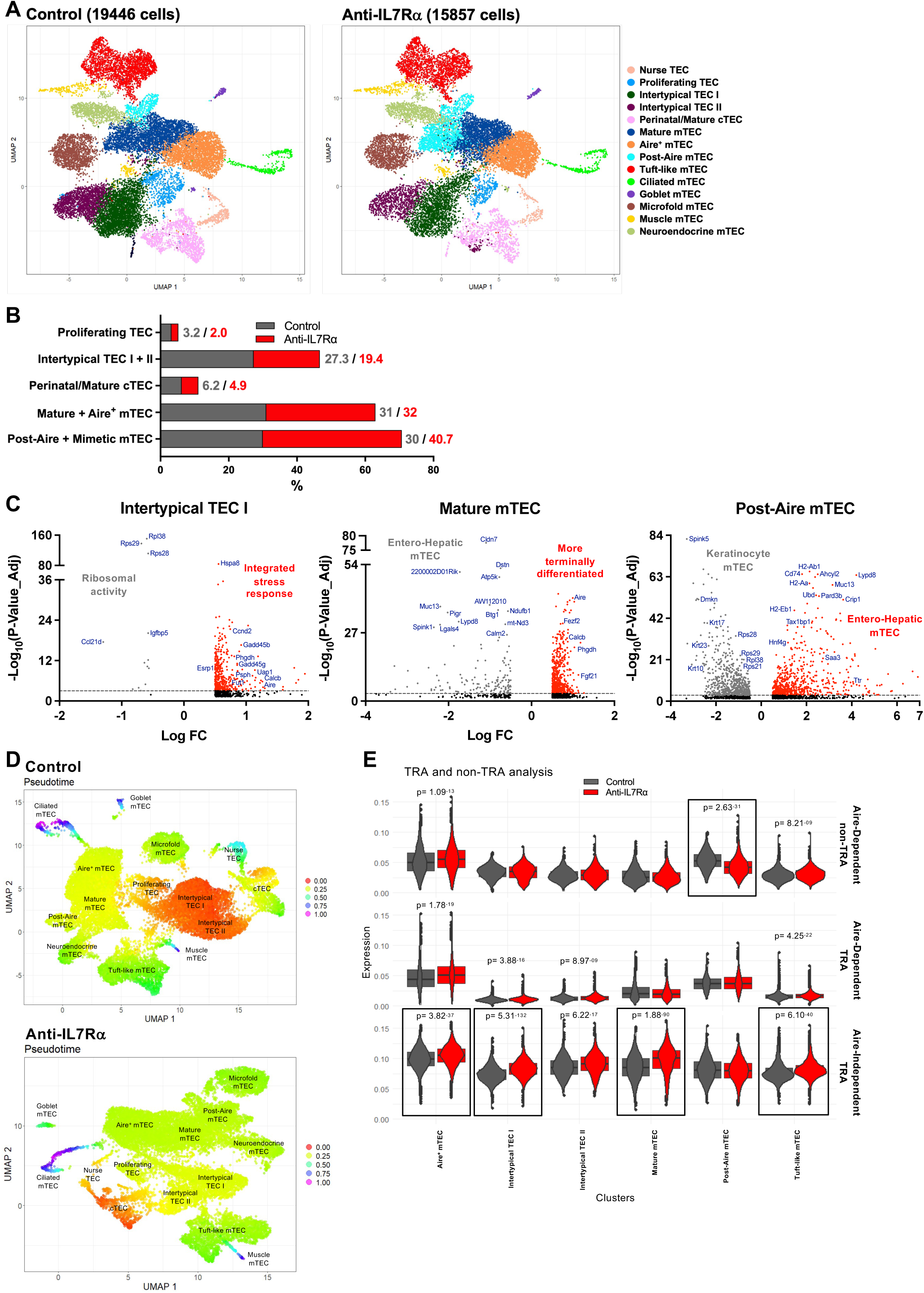
Altered TEC diversity and differentiation trajectory following first-wave TSP depletion. **(A)** UMAP visualization of single cell RNA sequencing of TEC from P30 control (left) and anti-IL7Rα (right) mice (19,446 control and 15,857 anti-IL7Rα cells, N=2 independent experiments per group). Fourteen clusters were defined and annotated based on canonical marker expression. Colors indicate distinct TEC subsets. **(B)** Frequencies of proliferating TEC, intertypical TEC I + II, perinatal/mature cTEC, mature + Aire⁺ mTEC, and post-Aire + mimetic mTEC. Mimetic mTEC include tuft-like, ciliated, goblet, microfold, muscle, and neuroendocrine subsets. **(C)** Volcano plots showing differentially expressed genes (DEG) between groups in intertypical TEC I (left), mature mTEC (middle), and post-Aire mTEC (right) subsets. **(D)** Pseudotime UMAP showing inferred TEC differentiation trajectories in control (top) and anti-IL7Rα (bottom) mice. Pseudotime values are shown as a continuous color scale (early = red, intermediate = yellow/green, late = blue/magenta). **(E)** Violin plots of Aire-dependent TRA, Aire-independent TRA, and Aire-dependent non-TRA across TEC populations (Aire⁺ mTEC, intertypical TEC I/II, mature mTEC, post-Aire mTEC, tuft-like mTEC). Black boxes highlight populations with significant differences (p values indicated).

Overall, cluster-defining genes were largely conserved between groups (**Supplementary Fig. 5A**; **Supplementary Table 2**). However, marked differences emerged within mature mTEC populations, including Aire⁺ and post-Aire subsets. In control mice, mature mTEC were defined by *Nos1*, *Aoc1*, and *Smtnl1*, whereas in anti-IL7Rα mice this cluster was characterized by *Prokr2*, *Resp18*, and *Gm10165*. Aire⁺ mTEC in controls expressed *Aire*, *Calcb*, and *Hdc*, while the corresponding cluster in anti-IL7Rα mice was instead defined by *Hal*, *Vmo1*, and *S100a8*, with *Aire* notably absent among the defining transcripts, indicating a loss of cluster-restricted *Aire* expression.

Post-Aire mTEC in control mice expressed *Spink5*, *Pdzk1ip1*, *Dmkn*, *Tacstd2*, and keratinocyte-associated transcripts^9^. In contrast, post-Aire mTEC from anti-IL7Rα mice were defined by entero-hepatic markers including *Pigr*, *Muc13*, and *Lgals4*, characteristic of mimetic TEC^9,25^ (Supplementary Fig. 5A; Supplementary Table 2). Moreover, four of the five defining post-Aire transcripts were shared with goblet and microfold TEC clusters in anti-IL7Rα mice, indicating blurred lineage boundaries. Consistently, hierarchical clustering revealed that post-Aire mTEC grouped with goblet TEC in controls but with microfold TEC in anti-IL7Rα mice. Proliferating TEC clustered with intertypical and cTEC/nurse TEC in controls, but instead associated with ciliated, Aire⁺, and mature mTEC subsets in anti-IL7Rα mice (**Supplementary Fig. 5B**).

Quantitatively, anti-IL7Rα mice exhibited reduced frequencies of proliferating TEC, intertypical TEC, and perinatal/mature cTEC, accompanied by a relative expansion of post-Aire and mimetic mTEC subsets (41% vs. 30% in controls; **Fig. 3B**; **Supplementary Fig. 5C**).

Notably, entero-hepatic post-Aire mTEC alone accounted for 10% of total TEC in anti-IL7Rα mice, compared with 2.3% in controls.

Together, these data demonstrate that depletion of first-wave TSP alters TEC differentiation trajectories, leading to mis-specified mature mTEC populations and an abnormal accumulation of post-Aire and mimetic mTEC subsets in the adult thymus.

### Accelerated maturation of mTEC in anti-IL7Rα mice

Differential gene expression (DEG) analyses across TEC subsets (**Fig. 3C**; **Supplementary Fig. 5D**; **Supplementary Table 3**) indicated five major alterations emerged in anti-IL7Rα mice: (i) a widespread downregulation of ribosomal genes (*Rps29*, *Rpl38*, *Rps28*) across most TEC subsets, with the exception of mature mTEC, accompanied by their upregulation in perinatal/mature cTEC; (ii) induction of stress-associated programs (*Gadd45b*, *Hspa8*, *Uap1*) in intertypical TEC I; (iii) ectopic upregulation of *Aire* and *Fezf2* in intertypical TEC I and mature mTEC; (iv) downregulation of enterohepatic mimetic genes (e.g., *Muc13*, *Lypd8*) in mature mTEC; and (v) a shift in post-Aire mTEC identity from a keratinocyte-like signature (*Krt10*, *Spink5*, *Dmkn*) in controls toward an entero-hepatic program (*Muc13*, *Lypd8*, *Hnf4γ*) in anti-IL7Rα mice^25^.

Together, these changes revealed broad transcriptional dysregulation across mTEC subsets, characterized by enhanced maturation signatures in intertypical and mature mTEC, expansion of entero-hepatic post-Aire mTEC, and evidence of an environmentally integrated stress response^26^, including reduced ribosomal gene expression and decreased translational activity. In contrast, perinatal/mature cTEC displayed increased ribosomal and proliferative activity (**Supplementary Fig. 5D**), potentially compensating for their reduced frequency and consistent with the failure of anti-IL7Rα mTEC to expand between P15 and P60 (**Fig. 1B**).

Pseudotime analysis further supported accelerated TEC differentiation in anti-IL7Rα mice. In controls, differentiation trajectories originated mainly from proliferating and intertypical TEC, with large distances separating progenitors from terminally differentiated subsets (**Fig. 3D**, **upper panel**). In contrast, anti-IL7Rα trajectories initiated from perinatal/mature cTEC and progressed rapidly through proliferating and intertypical states toward terminal fates, with reduced distances between intermediate and terminal subsets (**Fig. 3D**, **lower panel**). This was consistent with DEG data showing expression of mature and Aire-associated genes within intertypical TEC, indicating premature or dysregulated maturation.

Flow cytometric analysis of mTEChi subsets using CD24 and Sca-1, which discriminate late-Aire and post-Aire compartments^27^, corroborated these findings. Late-Aire CD24⁺Sca-1⁻ mTEC^hi^ were overrepresented in anti-IL7Rα mice from P15 to P60, and CD24⁺Sca-1⁺ post-Aire mTEC^hi^ were increased between P15 and P30, consistent with transcriptional and compositional analyses (**Supplementary Fig. 6**).

Cell-cycle scoring revealed similar global cycling dynamics across most TEC subsets, but proliferating TEC in anti-IL7Rα mice, despite reduced frequency, accumulated in S phase and showed enrichment of mitotic and chromosomal organization pathways (**Fig. 3B**; **Supplementary Fig. 7A–B**), indicating increased proliferative pressure. Gene ontology analysis further highlighted divergent functional identities of post-Aire mTEC, with enrichment of epidermal differentiation and antimicrobial pathways in controls, and enterohepatic programs in anti-IL7Rα mice (**Supplementary Fig. 7B**).

Finally, to assess functional consequences for promiscuous gene expression, we quantified Aire-dependent and Aire-independent TRA across TEC subsets (**Fig. 3E**; **Supplementary Table 4**). Intertypical TEC I and II from anti-IL7Rα mice expressed higher levels of Aire-independent, and to a lesser extent Aire-dependent, TRA, reflecting their advanced differentiation state. Mature mTEC showed a similar bias toward Aire-independent TRA. Aire⁺ and tuft-like mTEC expressed broad repertoires of TRA and non-TRA in both groups, whereas post-Aire mTEC differed markedly, consistent with their distinct cellular identities and antigen presentation profiles.

Collectively, these data demonstrate that depletion of the first wave of TSP accelerates TEC differentiation, biases terminal maturation toward entero-hepatic mimetic fates, and shifts TRA expression toward predominantly Aire-independent programs, thereby reshaping the epithelial landscape and potentially compromising central tolerance.

### The first wave of TSP impacts TEC regeneration after injury

Sublethal total-body irradiation (SL-TBI) at P60 induces a transient loss of thymocytes and TEC, which normally recover within 35 days (D) post-irradiation (PI) ^28^ (**Fig. 4A**). Before irradiation, thymocyte numbers were comparable between control and anti-IL7Rα mice, and both groups exhibited similar thymocyte depletion and recovery kinetics following SL-TBI (**Fig. 4B**).

**Figure 4.**
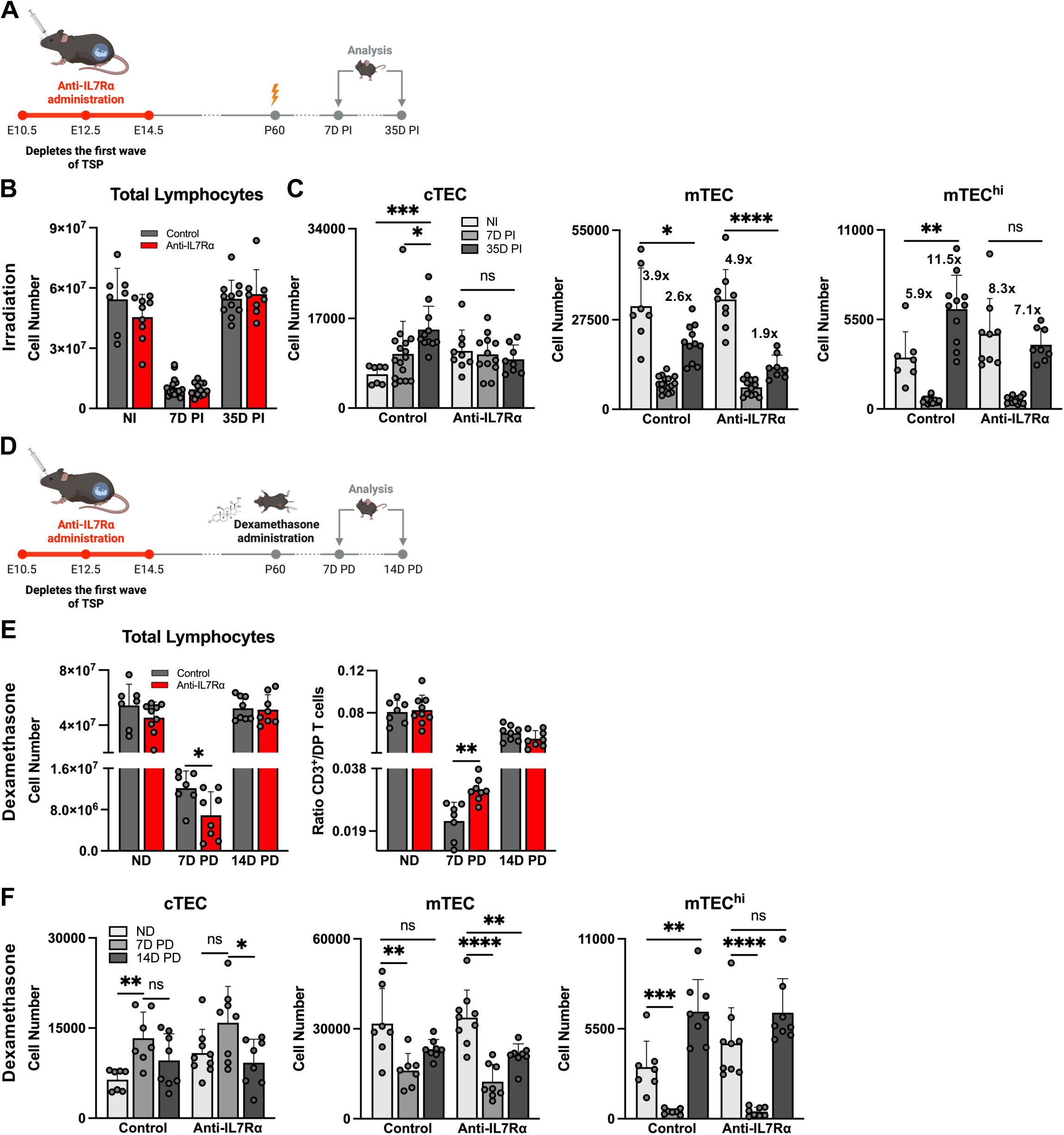
Impaired regenerative response of lymphocytes and TEC following injury in first wave TSP-depleted mice. **(A)** Experimental setup: P60 mice were sublethally irradiated (500 rad) and analyzed at 7- or 35-days (D) post-irradiation (PI). **(B)** Total lymphocyte numbers in non-irradiated (NI), 7D, and 35D PI mice. **(C)** Numbers of cTEC (left), mTEC (middle), and mTEC^hi^ mTEC (right) in NI, 7D, and 35D PI mice. The numbers above the bar graphs indicate the fold change between the two values. **(D)** Experimental setup: P60 mice were treated with dexamethasone (20 mg/kg) and analyzed at 7- or 14D post-dexamethasone (PD). **(E)** Total lymphocyte numbers (left) and ratio of CD3⁺ to double-positive (DP) T cells (right) in non-dexamethasone (ND), 7D, and 14D PD mice. **(F)** Numbers of cTEC (left), mTEC (middle), and mTEC^hi^ mTEC (right) in ND, 7D, and 14D PD mice. Each dot = individual mouse. Data are presented as the mean ± SD of at least 2 independent experiments. *p < 0.05, **p < 0.01, ***p < 0.001, ****p < 0.0001 (Unpaired T-test).

cTEC numbers in anti-IL7Rα mice failed to expand after SL-TBI (**Fig. 4C**, **left**) as observed in control mice and mTEC loss at 7D PI was more pronounced and recovery by 35D PI was substantially impaired compared with controls (**Fig. 4C**, **middle**). Between D7 and D35 PI, mTEC increased 1.9-fold in anti-IL7Rα mice versus 2.6-fold in controls. Although mTEC^hi^ cells showed partial recovery relative to total mTEC, they were more radiosensitive in anti-IL7Rα mice and failed to return to baseline levels by 35D PI (**Fig. 4C**, **right**), in contrast to control mice in which mTEC^hi^ numbers exceeded baseline by this time point.

We further treated P60 mice with dexamethasone, which induces thymic atrophy with faster recovery than irradiation^29^ (**Fig. 4D**). Both groups exhibited acute thymocyte loss; however, anti-IL7Rα mice showed a delayed rebound at 7D post-dexamethasone (PD), characterized by reduced total lymphocyte numbers and an increased proportion of mature CD3⁺ thymocytes relative to immature populations (**Fig. 4E**). In these mice, cTEC failed to expand at 7D PD, while mTEC were more severely affected and did not recover by 14D PD. Similarly, mTEC^hi^ numbers declined more markedly and failed to rebound after dexamethasone treatment (**Fig. 4F**).

Together, these data demonstrate that despite near-normal thymocyte and TEC numbers at steady state, mice lacking the full contribution of the first wave of TSP exhibit heightened sensitivity of both cTEC and mTEC to injury and a persistent defect in epithelial regeneration. These findings indicate that first-wave TSP or their progeny establish a long-lasting capacity for thymic resistance and efficient recovery following stress.

### The first wave of TSP prevents tolerance loss and autoimmunity in aged mice

We next assessed age-related defects in peripheral immune tolerance. Histopathological analysis at 9 months revealed prominent immune cell infiltrates in salivary and lacrimal glands, as well as in the colon, in anti-IL7Rα mice but not in control animals (**Fig. 5A–B**). We found a marked accumulation of CD4⁺ T cells in the colon of anti-IL7Rα mice, with frequencies and numbers increased by one to two orders of magnitude in 7 of 9 animals, compared with only 1 of 8 controls (**Fig. 5C**; **Supplementary Fig. 8**). This increase was associated with elevated numbers of CD44⁺ effector/memory CD4⁺ T cells and FOXP3⁺ RORγt⁻ Treg cells (**Fig. 5C**). The CD8⁺ T cell compartment, regardless of activation status, was comparable between groups (**Fig. 5C**; **Supplementary Fig. 8B**).

**Figure 5.**
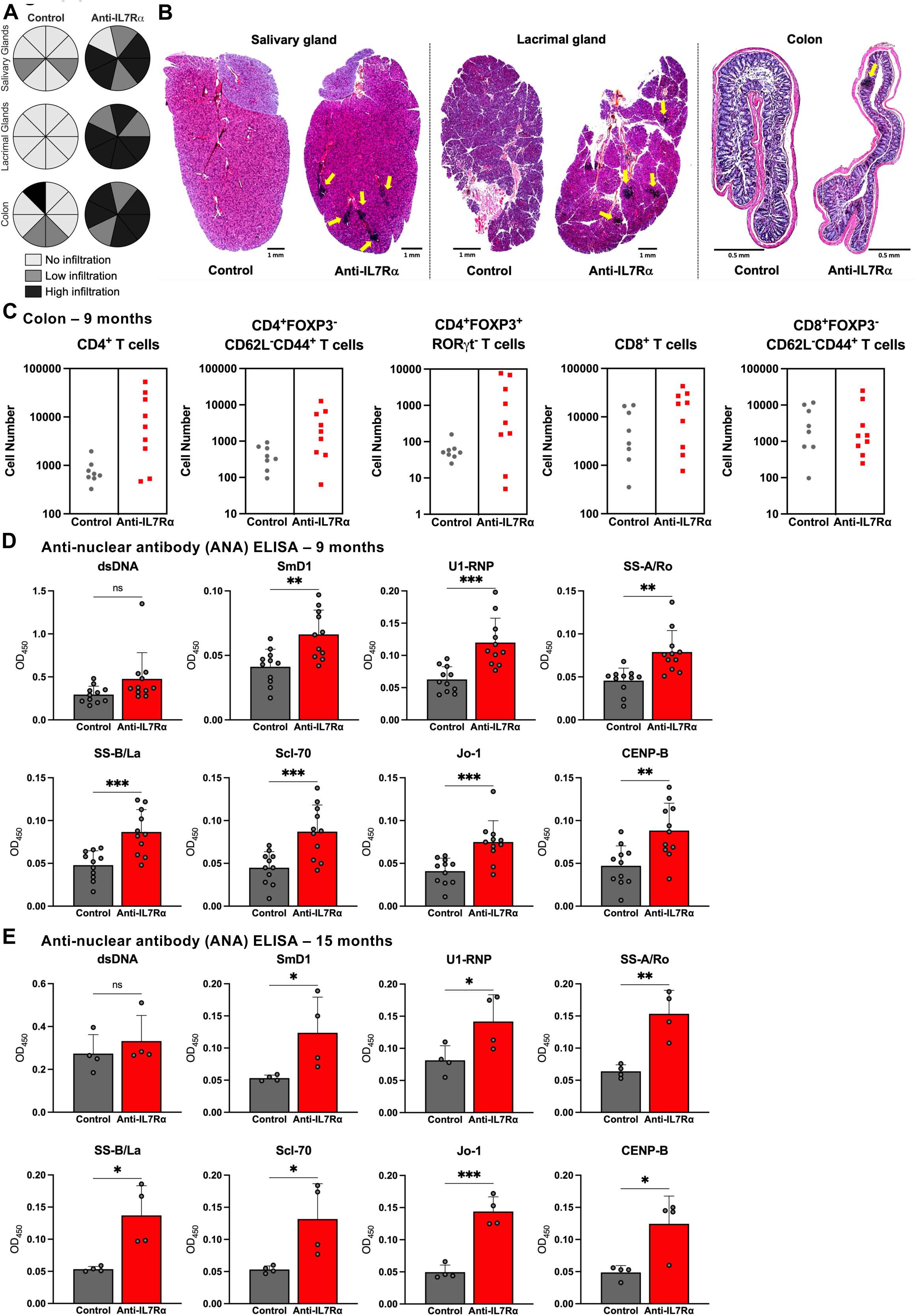
Depletion of first-wave TSP leads to impaired tolerance and autoimmunity. **(A)** Pie charts showing the degree of immune infiltration in the salivary/lacrimal glands and colon of 9-month-old mice: no infiltration (light grey), low (grey), and high (black). Each slice = one mouse. **(B)** Representative H&E images of infiltrates (yellow arrows) in salivary/lacrimal glands and colon at 9 months for both experimental groups. **(C)** Numbers of CD4⁺ (left), CD4⁺FOXP3^-^CD62L^-^CD44^+^ (middle left), CD4⁺FOXP3^+^RORψt^-^ (middle), CD8⁺ (middle right) and CD8^+^FOXP3^-^CD62L^-^CD44^+^ T (right) cells in colon of 9-month-old mice. Serum ANA detection at **(D)** 9 and **(E)** 15 months: dsDNA, SmD1, U1-RNP, SS-A/Ro, SS-B/La, Scl-70, Jo-1, and CENP-B. Data are presented as the mean ± SD of ≥ 2 independent experiments. ns, not significant. *p < 0.05, **p < 0.01, ***p < 0.001 (unpaired t test).

Further phenotypic analysis revealed increased numbers of FOXP3⁺ RORγt⁺ Treg cells and Th17-type CD4⁺ T cells in anti-IL7Rα mice (**Supplementary Fig. 8B**). Notably, a substantial fraction of infiltrating CD4⁺ and CD8⁺ T cells displayed a naïve phenotype (CD44⁻CD62L⁺) (**Supplementary Fig. 8B**), suggesting sustained recruitment of circulating T cells in response to local inflammation. Together, these findings indicate an ongoing autoreactive immune response accompanied by a compensatory accumulation of Treg cells that becomes evident with age.

To evaluate systemic autoimmunity, we quantified serum ANA at 9 and 15 months (**Fig. 5D–E**; **Supplementary Fig. 9**). Anti-IL7Rα mice exhibited significantly elevated levels of autoantibodies against multiple nuclear antigens (SmD1, U1-RNP, SS-A/Ro, SS-B/La, Scl-70, Jo-1, and CENP-B), whereas anti-dsDNA antibodies remained unchanged (**Fig. 5D–E**). ANA titers further increased with age in anti-IL7Rα mice, whereas they remained low and stable in controls (**Supplementary Fig. 9**).

Collectively, these data demonstrate that depletion of the first wave of TSP leads to a late-onset but severe breach of immune tolerance, characterized by tissue-specific inflammation and progressive autoreactivity. Thus, the progeny of the first wave of TSP exerts a long-lasting influence on thymic function and immune homeostasis well beyond the neonatal window of tolerance induction.

## Discussion

We show that early perturbation of thymic colonization leads to long-lasting alterations in TEC compartment composition, impaired regenerative capacity after injury, and a late-onset breakdown of immune tolerance.

Previous studies have established that first-wave TSP generate LTi and invariant γδ T cells, which are the first cells to produce RANK ligand, thereby initiating mTEC maturation and inducing Air^11,12,14^. Our findings extend this model by demonstrating that, despite the capacity of all mature T cells to produce RANK ligand, depletion of the first wave delays mTEC differentiation and durably imprints TEC organization and diversity. In normal mice, the thymus undergoes rapid expansion during the first 3–4 postnatal weeks, followed by progressive involution^16,30,31^. In contrast, TEC cellularity in anti-IL7Rα mice remained low and stable from P15 to P60, with a fragmented medulla composed of numerous small, poorly connected islets that persisted into adulthood^21^. The resulting elevated thymocyte-to-TEC ratio may reduce both the density and diversity of antigen presentation by mTEC, thereby impairing negative selection and Treg cell induction^32^. Notably, mice deficient in Ccl21a, LTβR, or LTα exhibit similar defects in medullary connectivity and three-dimensional organization^33–37^, and these molecules are produced by progeny of the first TSP wave^38,39^.

Consistent with an intact second wave of TSP, thymocyte cellularity recovered rapidly in anti-IL7Rα mice. However, the Treg cell compartment lagged behind conventional T cells and only normalized by P30. This delay is particularly relevant given the unique requirement for Aire-dependent tolerance during the perinatal period, when Aire⁺ mTEC promote the generation of a distinct wave of Foxp3⁺CD4⁺ Treg cells essential for lifelong self-tolerance^8,40^. Disruption of this developmental window, as observed in Aire-deficient mice, results in severe multiorgan autoimmunity^8,40^. Our data suggest that loss of first-wave TSP progeny selectively compromises perinatal Treg cell generation, contributing to the breakdown of tolerance observed in anti-IL7Rα mice.

While mTEC provide a unique repertoire of TRA, DC – particularly CD8α⁺ cDC1 – efficiently acquire and present Aire-regulated antigens^41–43^ from mTEC^4,41,44,45^, reinforcing central tolerance. The selective reduction of cDC1 observed at P30 suggests a shared, non-redundant developmental dependency between this DC subset and first-wave TSP progeny, further implicating early progenitors in the establishment of thymic antigen-presenting networks.

Single-cell transcriptomic analysis revealed that, in control mice, TEC differentiation proceeds through multiple routes originating from intertypical and proliferating TEC.

Intertypical TEC generate both Aire⁺ mTEC and terminal post-Aire populations^24,46^. Terminal mTEC identity is further shaped by “mimicry” programs, whereby mTEC adopt transcriptional features of peripheral tissues^9,10^. In contrast, TEC from anti-IL7Rα mice followed an altered trajectory originating from cTEC and progressing rapidly toward terminal differentiation, suggesting that loss of early thymocyte-derived cues accelerates or skews normal maturation. Consistently, Aire and Aire-associated transcripts (Calcb, Hdc), normally restricted to Aire⁺ mTEC, were aberrantly expressed in intertypical and mature mTEC subsets, indicative of premature or dysregulated maturation.

Recent work has shown that while most mimetic subsets emerge embryonically or early postnatally, keratinocyte and entero-hepatic mTEC arise later in life^47^, suggesting preferential shaping by thymocytes derived from the second TSP wave. The abrupt postnatal expansion of thymocytes in anti-IL7Rα mice may therefore drive abnormal TEC maturation, skewing the mimetic compartment toward entero-hepatic fates, exhausting progenitor pools, and inducing cellular stress responses. In line with this model, late- and post-Aire mTEC^hi^ subsets (CD24⁺Sca-1⁻ and CD24⁺Sca-1⁺)^27^ were increased from P15 to P60 in anti-IL7Rα mice. Together, these findings indicate that progressive, first-wave-driven maturation is essential for preserving TEC identity and maintaining balance among mimetic lineages.

Given that thymic architecture and function are established within a restricted embryonic window, preventing early developmental perturbations may be critical to ensure long-term immune competence. Identifying molecular mediators of first-wave imprinting – potentially involving RANKL, IL-22, IL-33, or related pathways^28,48,49^ – may therefore be insufficient to promote thymic repair.

In aged anti-IL7Rα mice, rising ANA titers between 9 and 15 months selectively involved autoantibodies associated with organ-specific diseases (SS-A/Ro, SS-B/La), characteristic of Sjögren’s syndrome^50–52^ linked to exocrine gland dysfunction and (Jo-1) linked to myositis and interstitial lung disease^53,54^, whereas anti-dsDNA antibodies linked to systemic lupus erythematosus^55–58^ remained unchanged. This serological profile, together with exocrine gland and colonic infiltration and preferential accumulation of Treg cells in tissues but not systemically, supports a breakdown of tissue-specific rather than systemic tolerance. The delayed onset of pathology suggests that early defects in TEC differentiation and antigen presentation progressively undermine central tolerance.

In summary, our study identifies the first wave of TSP as indispensable architects of thymic epithelial differentiation, regenerative capacity, and immune tolerance. By shaping TEC developmental trajectories and ensuring balanced mTEC subset composition, these progenitors establish a thymic microenvironment whose integrity cannot be restored later in life. Loss of first-wave TSP progeny therefore imprints a persistent epithelial vulnerability that ultimately manifests as selective failure of tissue-specific tolerance.

## Materials and Methods

### Mice

C57BL/6 mice were purchased from Envigo. Female mice (6–8 weeks old) and Rag2-GFP reporter mice were used for timed pregnancies. The day of vaginal plug detection was designated E0.5. All experiments were approved by the French Ministry of Education in accordance with institutional guidelines and European Directive 2010/63/EU.

### *In vivo* injections of anti-IL7Rα (A7R34) or anti-DCAT-1 antibodies

Pregnant females received intravenous injections of 1mg of anti–IL-7Rα antibody (clone A7R34^59^) at E10.5, E12.5, and E14.5. Control animals received an isotype-matched control antibody (DCAT-1). Offspring from treated females are referred to as anti-IL7Rα mice.

### Flow Cytometry

Single-cell suspensions were analyzed on an LSR Fortessa cytometer (BD Biosciences) and data processed using FlowJo. Antibodies used are listed in **Supplementary Table 1**. Dead cells were excluded using propidium iodide or fixable viability dyes. Intracellular FOXP3 staining was performed using a Foxp3 staining kit (eBioscience) according to the manufacturer’s instructions.

### Tissue processing

Spleen and thymic hematopoietic cells were obtained by mechanical dissociation. Peritoneal cavity cells were recovered as described^12^. Colonic lamina propria lymphocytes were isolated as described^60^. For TEC analysis, cell suspensions were obtained as described^12^.

### Thymic histology, imaging, and 3D reconstruction

Thymic lobes were fixed, embedded in agarose, and sectioned (100–150 μm). Sections were stained with DAPI, optically cleared, and imaged by confocal microscopy. 3D reconstruction was obtained as described^37,61^.

### Models of thymic damage

Thymic regeneration was assessed following sublethal total-body irradiation (500 rad) or a single intraperitoneal injection of 20 mg/kg of dexamethasone (Sigma-Aldrich). Thymi were analyzed at the indicated time points post-treatment.

### Histopathology

Salivary glands, lacrimal glands, and colon were fixed, paraffin-embedded, sectioned, and stained with hematoxylin and eosin. Lymphocytic infiltration was scored according to: No infiltration – absence of infiltrates in all sections (>2); Low infiltration – 1-2 infiltrates in all sections (>2). High infiltration – >2 infiltrates in all sections (>2).

### Anti-Nuclear Antibody detection

Serum autoantibodies were measured using a commercial mouse ANA ELISA (Signosis) that detects antibodies against eight nuclear antigens. Optical density was measured at 450 nm, and results were expressed as relative antibody levels.

### Single-Cell RNA-sequencing

Cortical and medullary TEC were enriched by depletion of CD45⁺ cells using magnetic beads, followed by flow cytometric sorting of EPCAM⁺CD45⁻ TEC subsets (Ly51⁺ cTEC and UEA-1⁺ mTEC). For each condition, 30,000 TEC were sorted per sample using an Aria III cytometer.

Single-cell libraries were generated using the Chromium X platform and the 10x Genomics 3′ Gene Expression v4 chemistry, following the manufacturer’s protocol. Libraries were quality-controlled, pooled at equimolar ratios, and sequenced on an Illumina NextSeq2000 instrument.

#### Read processing and quality control

Sequencing data were processed using Cell Ranger^62^ against the mouse GRCm39 reference genome. Downstream analyses were performed using Scanpy (Python)^63^. Low-quality cells were excluded based on gene complexity, mitochondrial content, and doublet detection^64^. Genes detected in fewer than 20 cells were removed. Data were normalized, log-transformed, and highly variable genes were selected per batch.

#### Dimensionality reduction, integration, and clustering

Principal component analysis was performed on scaled highly variable genes. Batch effects were corrected using Harmony^65^, and integrated embeddings were used to compute nearest-neighbor graphs for UMAP visualization and Leiden clustering^66^. Cluster annotation was based on differentially expressed marker genes^63^ and comparison with published TEC reference datasets^9,24,67^.

#### Cell-cycle analysis

Cell-cycle phase scores were calculated using established S-phase and G2/M gene signatures to assess proliferative states across TEC clusters^68^.

#### Differential expression and enrichment analyses

For sample-level comparisons, counts were aggregated into pseudobulk profiles^69^ and differential expression analysis was performed using DESeq2^70^. Genes meeting significance and fold-change thresholds were considered differentially expressed. Gene set enrichment analyses were conducted using curated GO, KEGG, and MSigDB gene sets^71^.

#### Pseudotime analysis

Cell-state trajectories were inferred using diffusion maps and diffusion pseudotime^72^, with root cells selected from progenitor-enriched TEC clusters based on marker expression.

## Statistical analysis

Data are presented as mean ± SEM or SD, as indicated. Statistical analyses were performed using GraphPad Prism, with tests specified in figure legends.

## Acknowledgments

We thank the Center for Translational Science (CRT) – Cytometry and Biomarkers Unit of Technology and Service (CB UTechS) at the Institut Pasteur for their support throughout this study, in particular S. Novault, S. Megharba, and S. Schmutz. We are grateful to the staff of the Institut Pasteur animal facility for exemplary mouse care, and to the Histopathology Core Facility for their assistance, especially D. Hardy, D. Hing, S. Loulizi, and M. Tichit. We also acknowledge F. Jagorel from the Biomics Platform, C2RT, Institut Pasteur, Paris, supported by France Génomique (ANR-10-INBS-09) and IBISA, for sequencing support. We warmly thank Marie-Pierre Mailhe for her technical assistance, and Paulo Vieira and all members of the Lymphocytes and Immunity Unit for their insightful discussions and continuous scientific input.

## Author contributions

G.N., A.C., F.S.S., A.B., N.L.A., P.P., A.C. designed the experiments; G.N., A.B., P.P., N.L.A., A.C. co-wrote the manuscript; G.N., A.C., F.S.S., C.C., A.G., A.B., P.F., A.C. performed experiments; and all authors contributed to the manuscript.

---

**Supplementary Figure 1.**
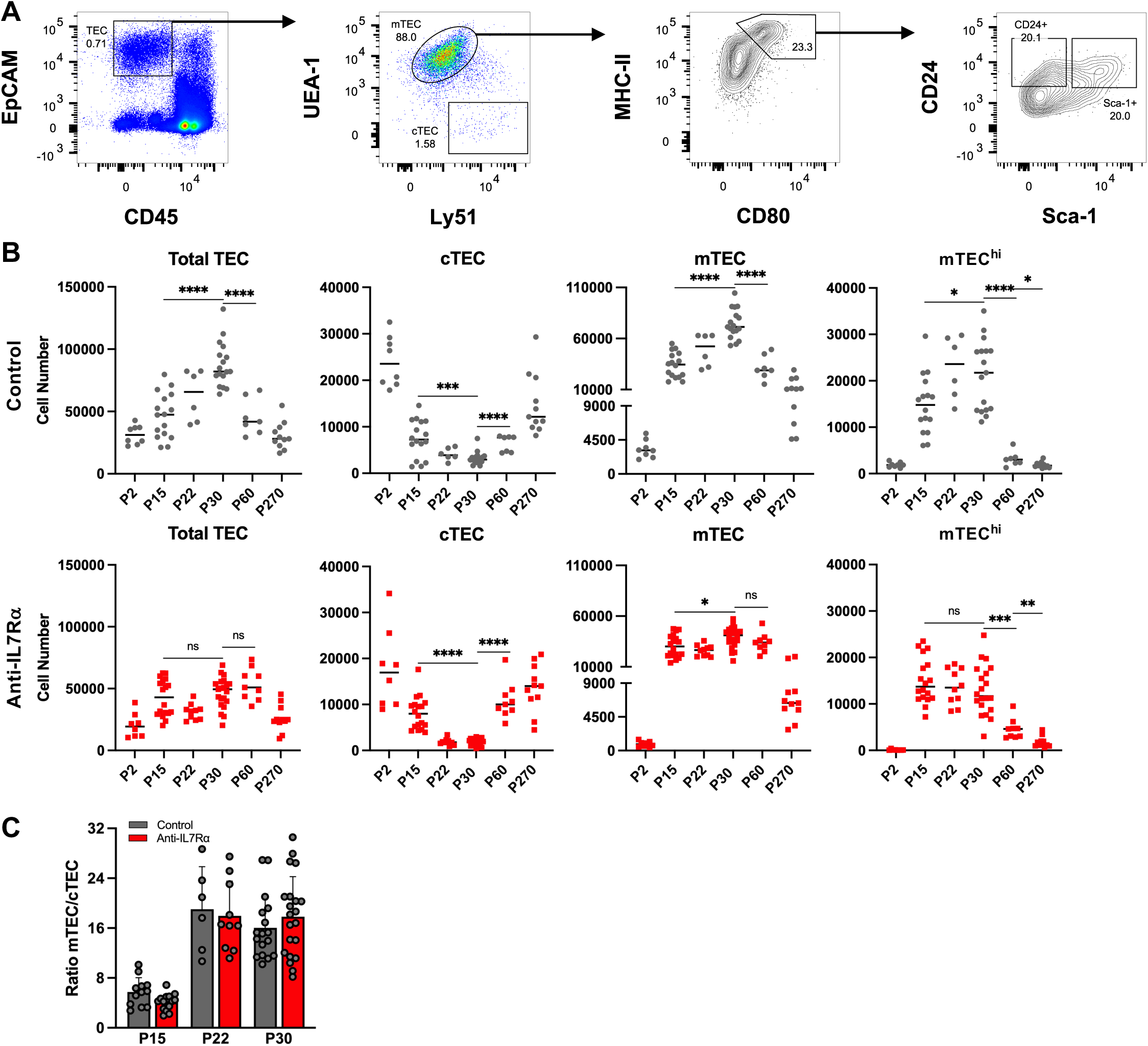
Postnatal alterations in TEC following first-wave TSP depletion. **(A)** Gating strategy used for TEC analysis. **(B)** Absolute numbers of total TEC (left), cTEC (middle left), mTEC (middle right), and mTEC^hi^ mTEC (right) at P2, P15, P22, P30, P60, and P270 in control and anti-IL7Rα mice. **(C)** Ratio of mTEC to cTEC at P15, P22, and P30 in both experimental groups. Each dot represents one thymus. Data are presented as the mean ± SD of ≥ 2 independent experiments. *p < 0.05, **p < 0.01, ***p < 0.001, ****p < 0.0001 (unpaired t test).

**Supplementary Figure 2.**
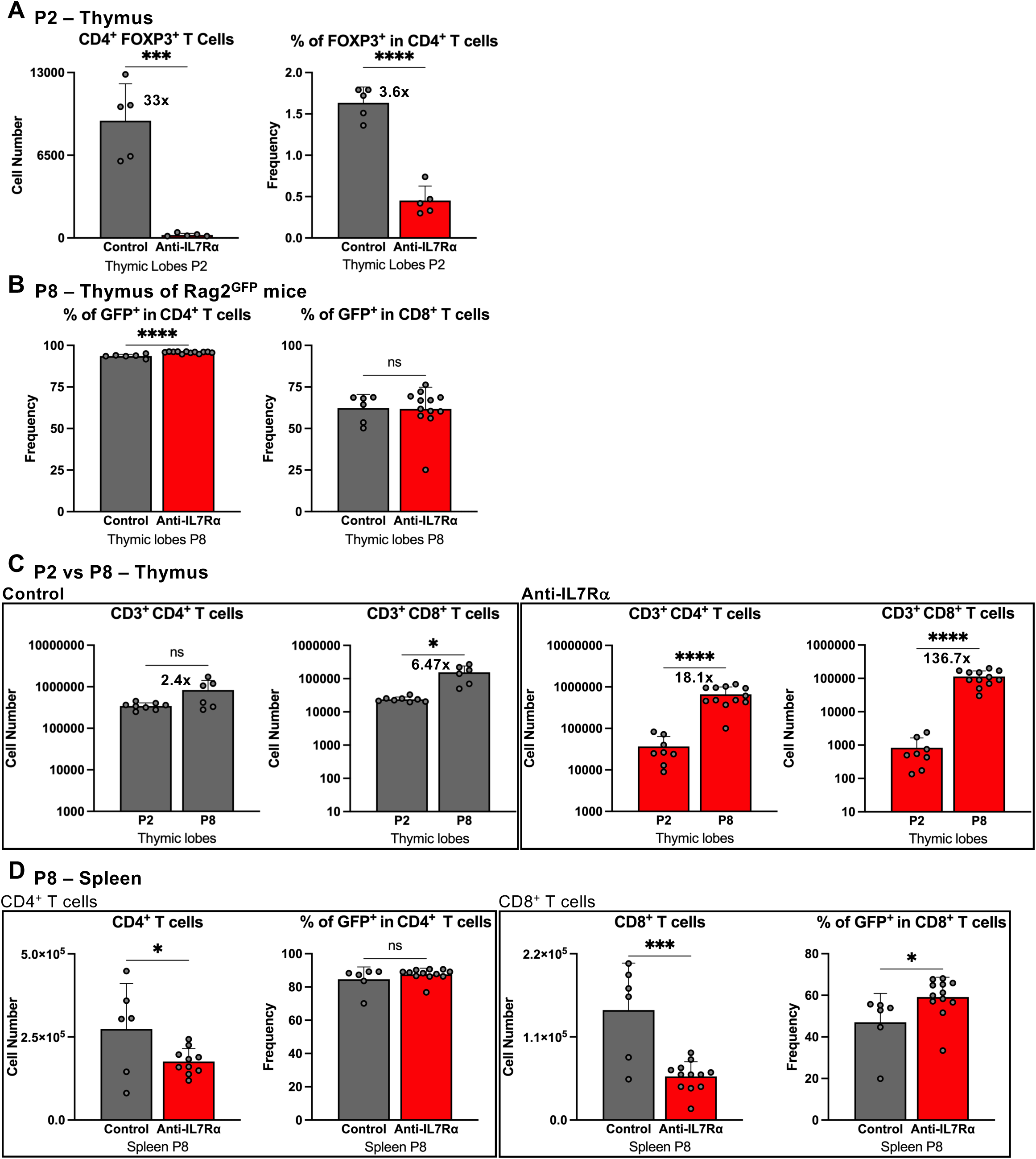
Sequential recovery of thymic and peripheral lymphocyte compartments following first-wave TSP depletion. **(A**) Absolute numbers (left), and frequencies (right) of CD4⁺FOXP3⁺ T cells in the thymus of P2 mice. The numbers above the bar graphs represent the fold change between the values of the two experimental groups. **(B)** Frequencies of GFP^+^ in CD3⁺CD4⁺ (left) and CD3⁺CD8⁺ (right) T cells in the thymus of P8 mice. **(C)** Comparison between the numbers of CD3^+^ CD4^+^ and CD8^+^ T cells at P2 and P8 in control (left)and anti-IL7Rα (right) mice. The numbers above the bar graphs show the fold change between the two time points. **(D)** Absolute numbers of CD4^+^ and frequencies of GFP^+^ in CD4^+^ T cells (left) and absolute numbers of CD8^+^ and frequencies of GFP^+^ in CD8^+^ T cells (right) in the spleen of P8 mice. Data are presented as the mean ± SD of ≥ 2 independent experiments. *p < 0.05, **p < 0.01, ***p < 0.001, ****p < 0.0001 (unpaired t test).

**Supplementary Figure 3.**
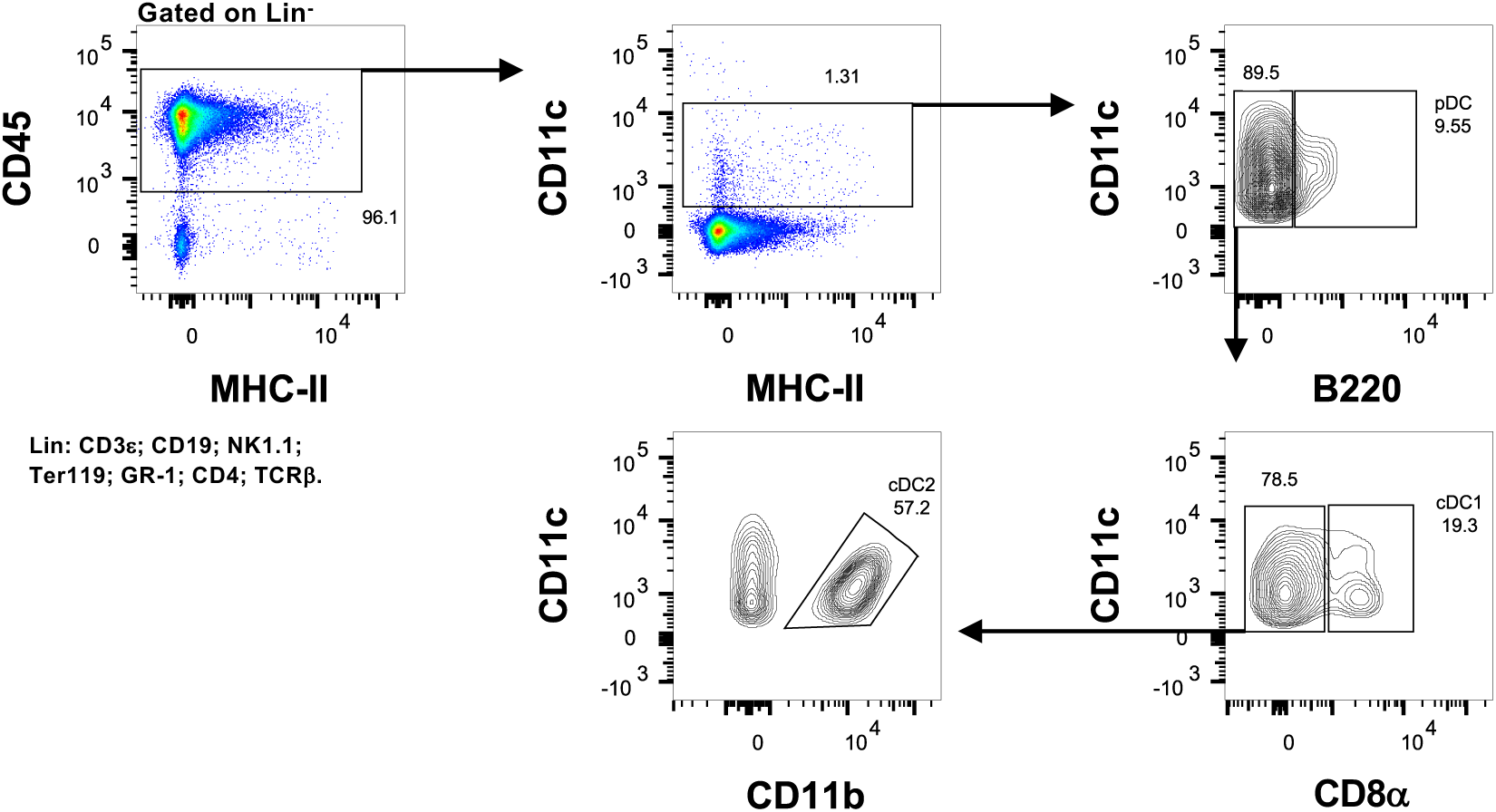
Gating strategy for thymic dendritic cell subset analysis. Representative flow cytometry plots illustrating the sequential gating strategy used to identify and discriminate the different dendritic cell (DC) subsets in the thymus.

**Supplementary Figure 4.**
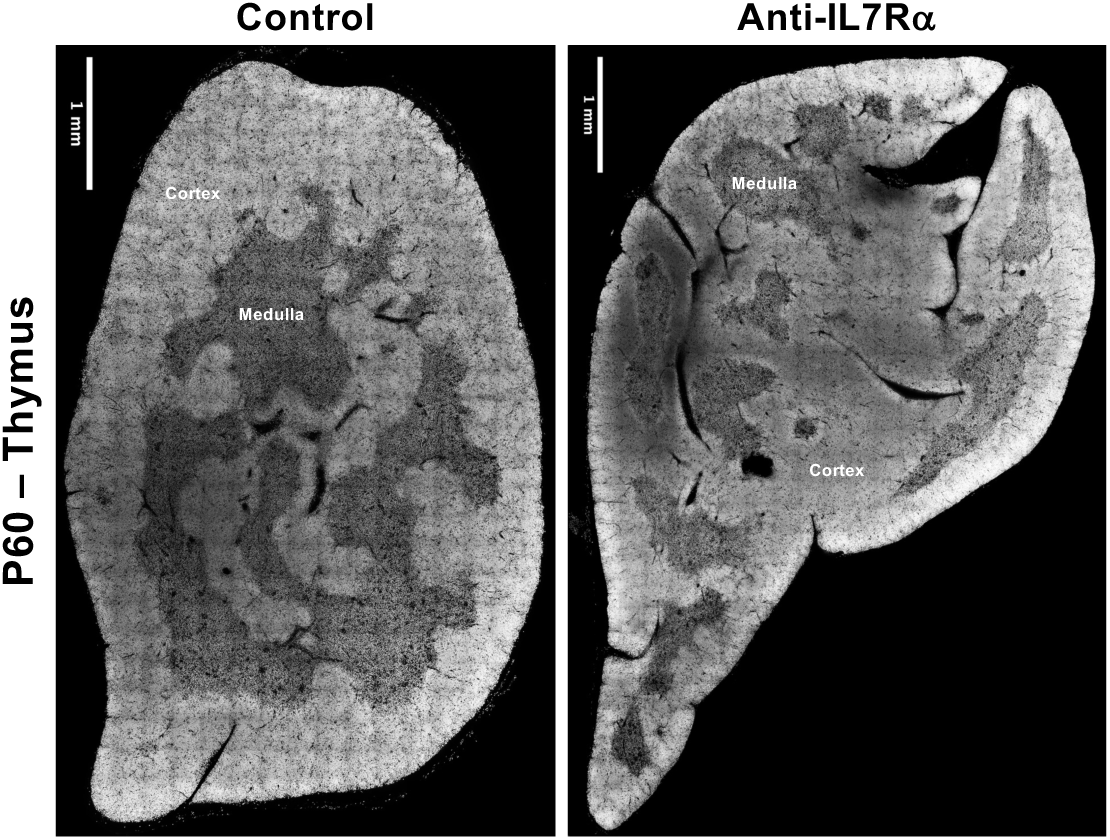
Persistent abnormalities in medullary compartment organization in animals with reduced first-wave TSP. Representative DAPI-stained thymic sections at P60 in control and anti-IL7Rα mice.

**Supplementary Figure 5.**
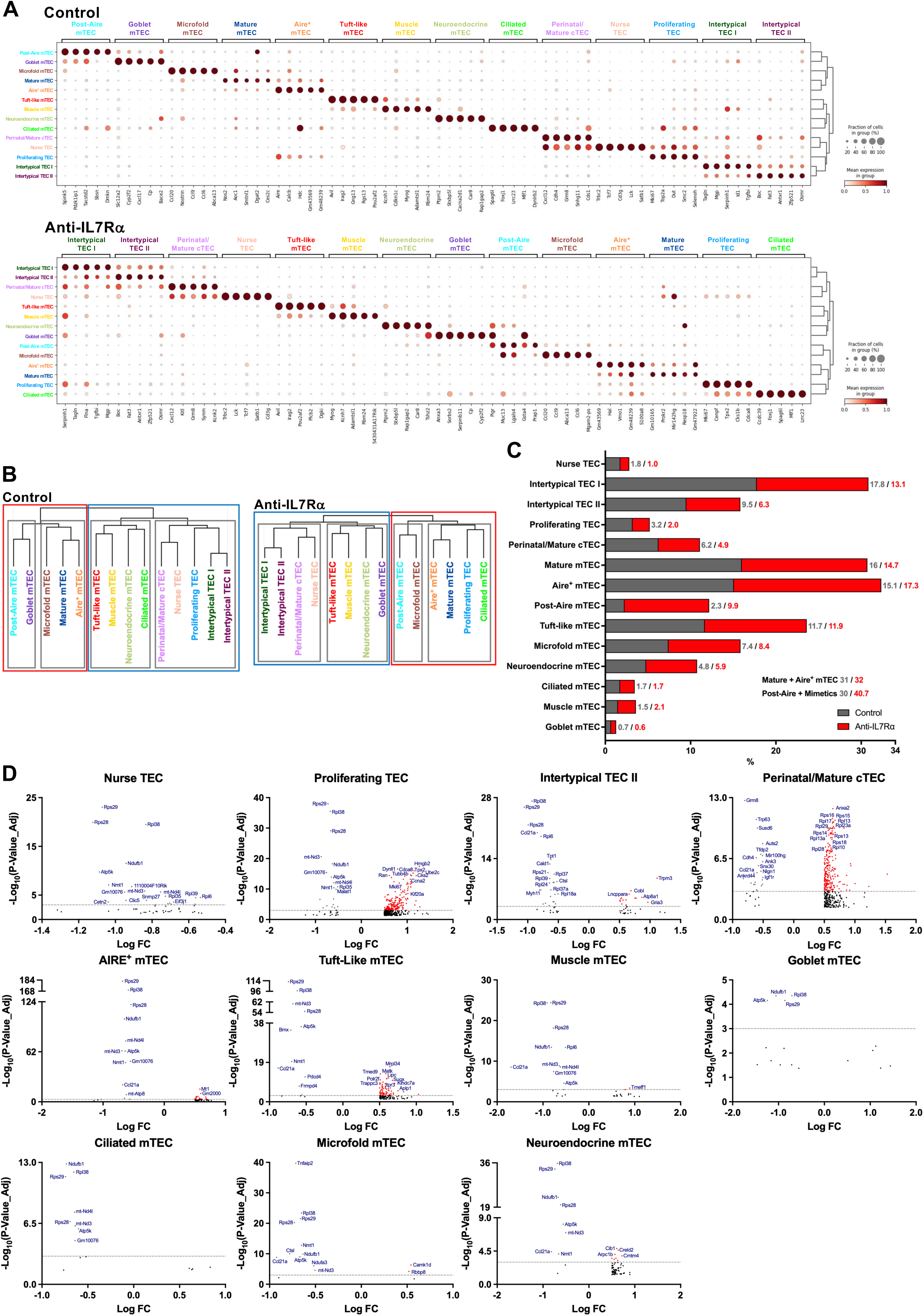
Molecular and functional profiling of TEC subsets after first-wave TSP depletion. **(A)** Marker gene expression profiles used for TEC cluster definition in single-cell transcriptomic analysis data sets in control (top) and anti-IL7Rα (bottom) thymi. Each dot indicates the fraction of cells within a cluster expressing a given marker, with dot size representing the proportion of positive cells and color intensity reflecting the average expression level. **(B)** Hierarchical tree illustrating the developmental relationships and relative distances between TEC clusters in control (left) and anti-IL7Rα (right) mice. **(C)** Relative frequency of TEC subsets across experimental groups. **(D)** Volcano plots showing differentially expressed genes (DEG) between groups in nurse TEC, proliferating TEC, intertypical TEC II, perinatal/mature cTEC, Aire^+^ mTEC, tuft-like mTEC, muscle mTEC, goblet mTEC, ciliated mTEC, microfold mTEC and neuroendocrine mTEC.

**Supplementary Figure 6.**
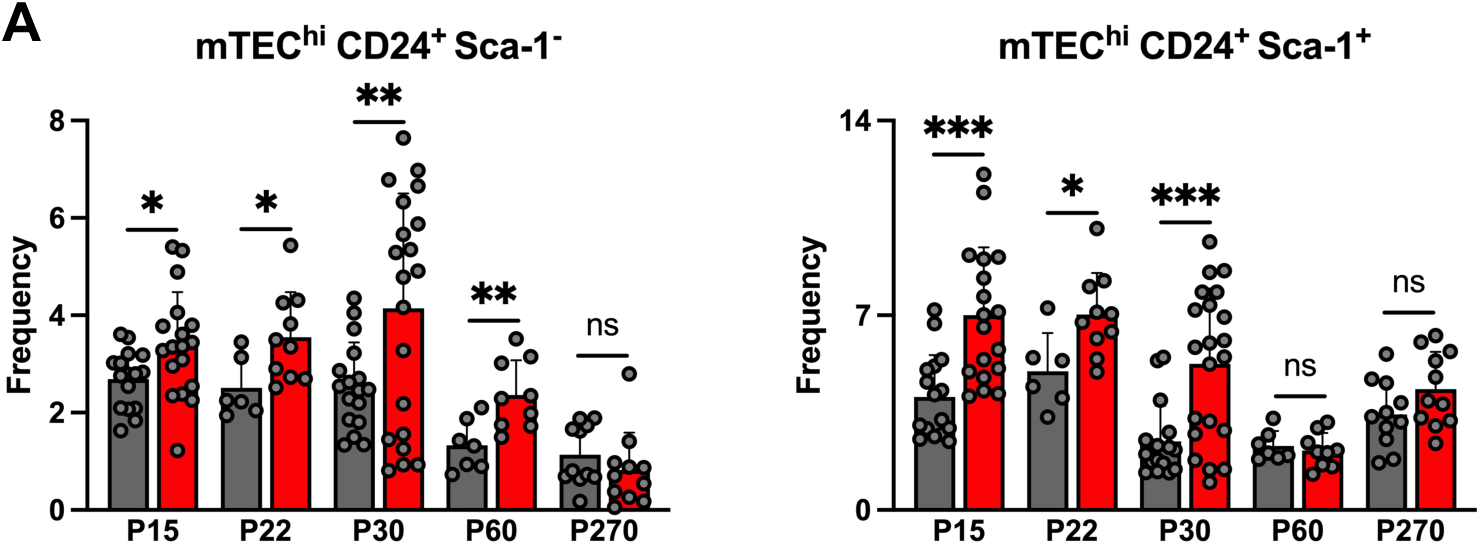
Long-lasting increase in mTEC maturation in animals with reduced first-wave TSP. Frequency of mTEC^hi^ CD24^+^ Sca-1^-^ (left) and CD24^+^ Sca-1^+^ (right) between P15 and P270.

**Supplementary Figure 7.**
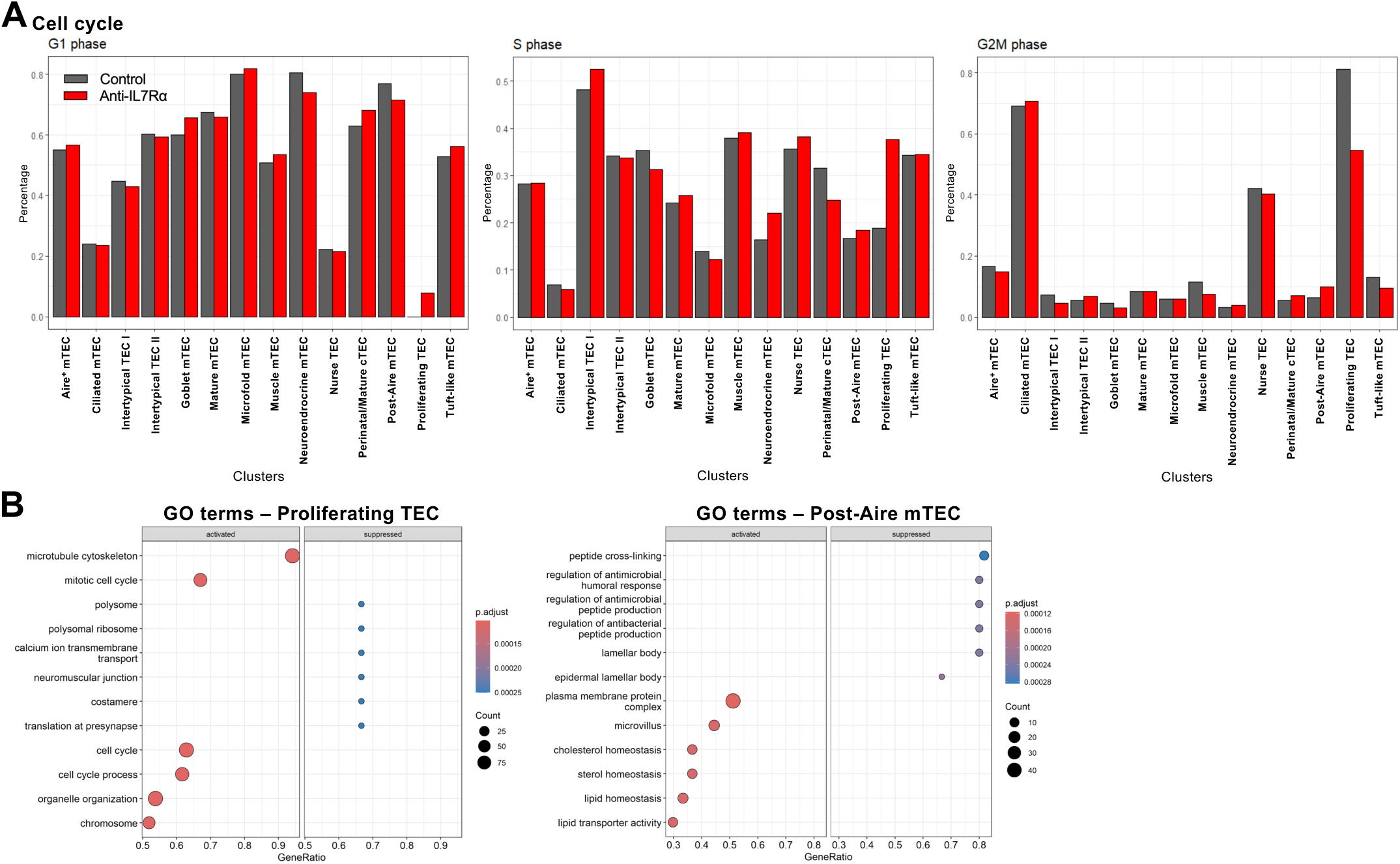
Altered cell cycle dynamics and transcriptional programs in TEC subsets of anti-IL7Rα mice. **(A)** Distribution of cells from each TEC cluster across cell cycle phases (G1, S, G2/M). **(B)** Gene ontology (GO) enrichment analysis showing pathways significantly upregulated or downregulated in proliferating TEC (left) and post-Aire mTEC (right) subsets in anti-IL7Rα compared with control mice.

**Supplementary Figure 8.**
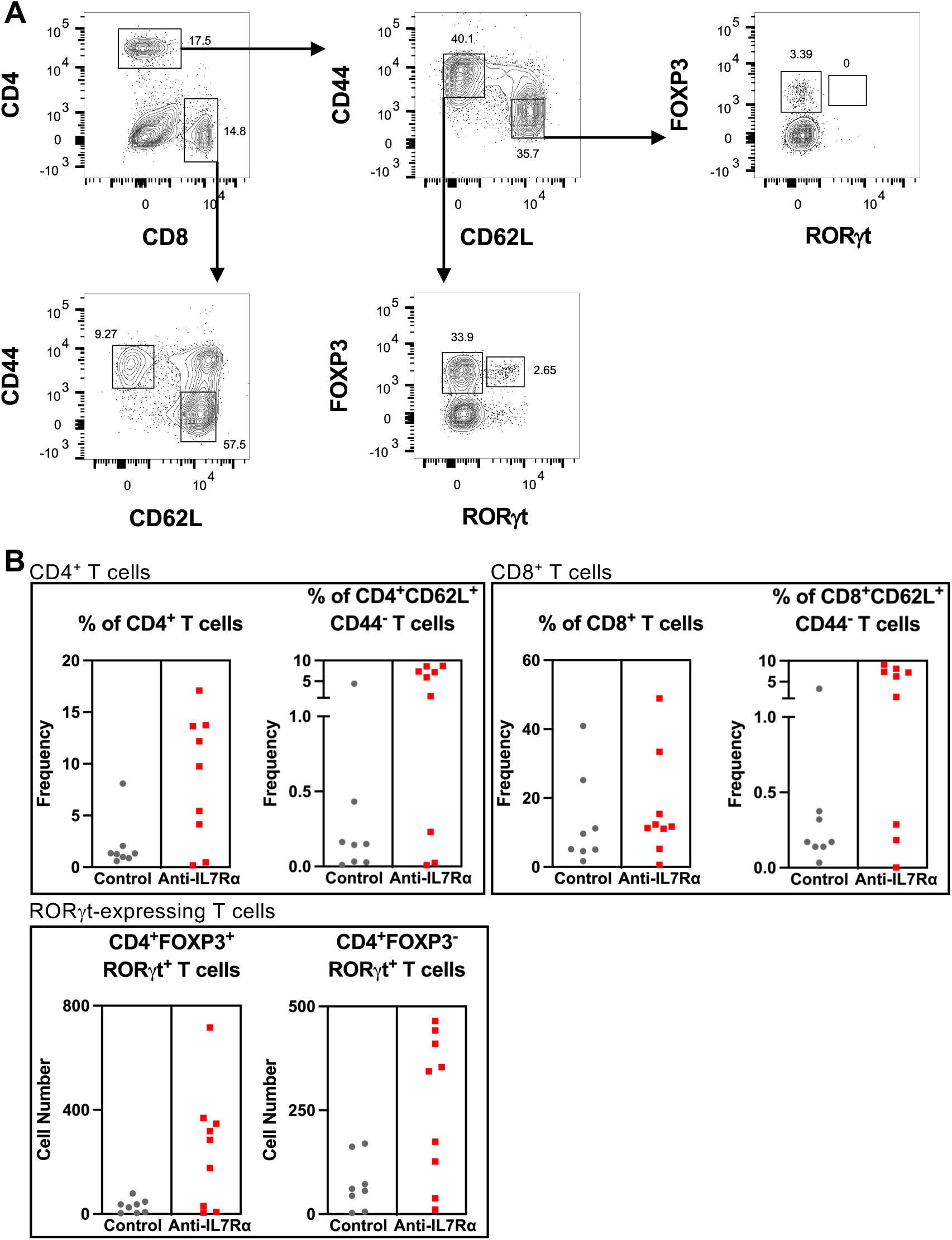
Altered colon T cell composition in 9-month-old anti-IL7Rα mice. **(A)** Flow cytometry gating strategy for the analysis of colonic T cells. Initial gating on CD4 vs CD8 distinguishes single-positive subsets. CD4⁺ and CD8⁺ T cells were first identified and further analyzed for CD44 and CD62L expression to distinguish naïve (CD44⁻CD62L⁺) and effector/memory (CD44⁺CD62L⁻) subsets. Within the CD4⁺ compartment, FOXP3 and RORγt expression was assessed to discriminate FOXP3⁺RORγt⁻ Treg cells from Th17-type RORγt⁺ Treg cells. **(B)** Frequency of CD4^+^ and of CD4^+^CD62L^+^CD44^-^ T cells (upper left), frequency of CD8^+^ and of CD8^+^CD62L^+^CD44^-^ T cells (upper right), absolute numbers of CD4^+^FOXP3^+^RoRψt^+^ (bottom left) and of CD4^+^FoxP3^-^ RoRψt^+^ (bottom right) colonic T cells in control and anti-IL7Rα mice at 9 months of age.

**Supplementary Figure 9.**
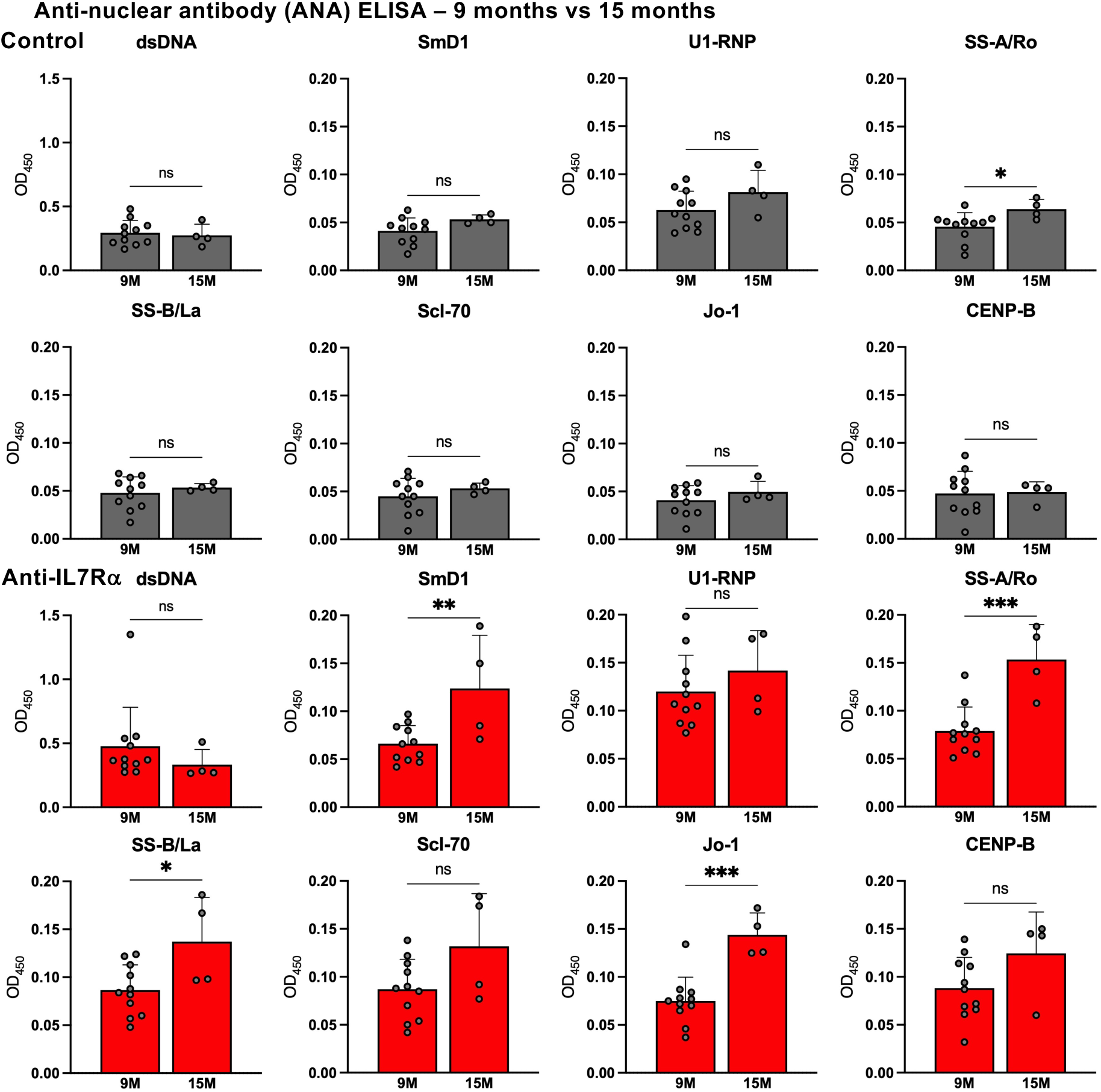
Progressive increase in serum ANA titers in anti-IL7Rα mice with age. Comparative analysis of ANA titers between 9- and 15-month-old mice in each experimental group.

**Supplementary Table 1.** List of antibodies used for flow cytometry.

| Protein | Conjugation | Clone | Catalog # | Manufacturer | Species | Used for the detection of |
| --- | --- | --- | --- | --- | --- | --- |
| B220 | FITC | RA3-6B2 | 553088 | BD Biosciences | Rat | Dendritic cells |
| EpCAM | BV421 | G8.8 | 563214 | BD Biosciences | Rat | Thymic Epithelial Cells |
| UEA-1 | Biotin | - | B-1065 | Vector Laboratories | Rat | Thymic Epithelial Cells |
| CD80 | FITC | 16-10A1 | 11080182 | BD Biosciences | Rat | Thymic Epithelial Cells |
| Ly51 | PE | BP-1 | 553735 | BD Biosciences | Rat | Thymic Epithelial Cells |
| CCR6 | PB | 140706 | 564736 | BD Biosciences | Rat | T cells |
| CD11b | PB | M1/70 | 1106120 | SONY | Rat | Dendritic Cells |
| CD11c | PE | N418 | 117308 | BD Biosciences | Rat | Dendritic Cells |
| MHC-II | BV711 | M5/114.15.2 | 563414 | BD Biosciences | Rat | Antigen-Presenting Cells |
| CCR6 | APC | 29-2L17 | 1249070 | SONY | Rat | T cells |
| FOXP3 | APC | FJK-16s | 17-5773-82 | BD Biosciences | Rat | T cells |
| FOXP3 | PE | 150D | 320007 | BD Biosciences | Rat | T cells |
| CD8a | PerCP-Cy5.5 | 53-6.7 | 551162 | BD Biosciences | Rat | T cells |
| CD3e | PE-Cy7 | 145-2C11 | 552774 | BD Biosciences | Rat | T cells |
| CD3e | Biotin | 145-2C11 | 1101520 | SONY | Rat | Lineage Depletion |
| CD3e | BV510 | 145-2C11 | 563024 | BD Biosciences | Rat | T cells |
| CD3e | BUV395 | 145-2C11 | 563565 | BD Biosciences | Rat | T cells |
| CD4 | Biotin | GK1.5 | 1102020 | SONY | Rat | Lineage Depletion |
| CD4 | BV786 | RM4.5 | 563727 | BD Biosciences | Rat | T cells |
| CD45 | BUV395 | 30-F11 | 564279 | BD Biosciences | Rat | Hematopoietic Cells |
| CD45 | PERCP-CY5.5 | 30-F11 | 1115660 | SONY | Rat | Hematopoietic Cells |
| CD62L | BUV395 | MEL-14 | 569400 | BD Biosciences | Rat | T cells |
| TCRb | Biotin | H57-597 | 1146020 | SONY | Rat | Lineage Depletion |
| CD8a | APC | 53-6.7 | 553035 | BD Biosciences | Rat | T cells |
| CD8a | PerCP-CY5.5 | 53-6.7 | 551162 | BD Biosciences | Rat | T cells |
| Gr-1 | Biotin | RB6-8C5 | 553125 | BD Biosciences | Rat | Lineage Depletion |
| NK1.1 | Biotin | PK136 | 1143520 | SONY | Rat | Lineage Depletion |
| CD69 | PE-Cy7 | H1.2F3 | B222011 | BD Biosciences | Rat | T cells |
| Ter119 | Biotin | TER-119 | 1181020 | SONY | Rat | Lineage Depletion |
| CD44 | BV510 | IM7 | 563114 | BD Biosciences | Rat | Thymocytes |
| TCRβ | BUV737 | H57-597 | 612821 | BD Biosciences | Rat | Thymocytes |
| TCRγδ | PERCP-CY5.5 | GL3 | 1190590 | SONY | Rat | T cells |
| RORγT | PE-CF594 | Q31-378 | 562684 | BD Biosciences | Rat | T cells |
| B220 | BV605 | RA3-6B2 | 1116215 | SONY | Rat | B cells |
| CD8a | PB | 53-6.7 | 558106 | BD Biosciences | Rat | T cells |
| CD5 | PE | 53-7.3 | 100608 | SONY | Rat | B cells |
| CD19 | APC-Cy7 | 1D3 | 551655 | BD Biosciences | Rat | B cells |
| CD19 | Biotin | 6D5 | 1177520 | SONY | Rat | Lineage Depletion |
| Streptavidin | APC | - | 2626035 | SONY | - |  |
| Streptavidin | BV650 | - | 563855 | BD Biosciences | - |  |
| Streptavidin | PB | - | S11222 | Invitrogen | - |  |
| Streptavidin | PE-Cy7 | - | 2626030 | SONY | - |  |
| Streptavidin | BUV737 | - | 564293 | BD Biosciences | - |  |

**Supplementary Table 2.** Marker genes defining each TEC subtype in control and anti-IL7Rα mice.

| Cluster | Aire-Dependent<br>TRA_pvalue | Aire-Dependent<br>nonTRA_pval | Aire-Independent<br>TRA_pval |
| --- | --- | --- | --- |
| Mature mTEC | 0.04 | 0.55 | 1.88<sup>-90</sup> |
| Post-Aire mTEC | 1 | 2.63<sup>-31</sup> | 1 |
| Nurse TEC | 0.87 | 1 | 0.19 |
| Microfold TEC | 0.26 | 0.86 | 0.001 |
| Aire<sup>+</sup> mTEC | 1.78<sup>-19</sup> | 1.09<sup>-13</sup> | 3.82<sup>-37</sup> |
| Ciliated mTEC | 0.01 | 1 | 8.66<sup>-07</sup> |
| Intertypical TEC II | 8.97<sup>-09</sup> | 0.31 | 6.22<sup>-17</sup> |
| Perinatal Mature cTEC | 1 | 0.01 | 1 |
| Intertypical TEC I | 3.88<sup>-16</sup> | 1 | 5.31<sup>-132</sup> |
| Tuft-like mTEC | 4.25<sup>-22</sup> | 8.21E-09 | 6.10<sup>-40</sup> |
| Neuroendocrine mTEC | 5.79<sup>-07</sup> | 0.002 | 2.28<sup>-13</sup> |
| Proliferating TEC | 1 | 1 | 1 |
| Goblet mTEC | 0.01 | 0.01 | 1 |
| Muscle mTEC | 0.007 | 0.15 | 0.02 |

**Supplementary Table 3.** Differentially expressed genes identified in each TEC cluster when comparing control and anti-IL7Rα mice.

**Supplementary Table 4.** Comparative analysis of Aire-dependent and Aire-independent tissue-restricted and non-tissue-restricted antigen expression across TEC subsets in control and anti-IL7Rα mice.

**Supplementary Video 1. 3D reconstruction of a thymic lobe from a control mouse.**

**Supplementary Video 2. 3D reconstruction of a thymic lobe from an anti-IL7Rα mouse.**

